# Biological and technical spatial variation in spatial transcriptomics

**DOI:** 10.64898/2026.10.13.758896

**Authors:** Kayla Jackson, Gennady Gorin, Lior Pachter

## Abstract

Spatial transcriptomics technologies enable the identification of spatially variable genes that can shed light on tissue structure and function. However, spatial variability of gene expression can be due not only to biological function but also technical artifact. We develop a model for spatially variable capture that forms part of a bursty transcription model, allowing us to distinguish biological variability from spatially variable technical noise. Our approach is implemented in a program called stipple, which we demonstrate can be useful for identifying and interpreting spatial gene expression variation from spatial transcriptomics assays.

## Introduction

Spatially-resolved transcriptomics (SRT) technologies measure gene expression while preserving the contextual information contained in intact tissues (Moses and Pachter, 2022). This allows for assignment of transcriptomic measurements to morphologically or functionally defined domains that are known to exist in many tissues. These domains are identifiable with histopathological techniques, but the approaches reveal limited information about the molecular programs that distinguish domains. Conventional single cell RNA sequencing (scRNA-seq) can measure these programs directly, but tissue dissociation precludes comparison of gene expression across histological features unless they are accurately dissected beforehand (Nichterwitz et al., 2016; Merritt et al., 2020). Additionally, there are gene expression signatures that do not respect the boundaries of any partition of the tissue, and thus cannot be recovered using methods that require dissociation.

Identification of genes with spatially distinct expression patterns is a primary focus of SRT data analysis. Genes that distinguish domains are spatially patterned by construction, since the pattern follows from the arrangement of the domains themselves. We refer to genes whose spatial variation is attributable to such discrete organization as domain-variable genes (DVGs). In contrast, gene expression can also vary across space due to processes that are not explained by discrete organization, including from expression gradients and local signaling. We use the term spatially-variable genes (SVGs) to describe this pattern of spatial variation.

Although both gene classes display spatial autocorrelation, they represent different biological phenomena. The first case reflects genes that are useful for characterizing features that define known domains. The second class captures processes that are not organized around any partition of the tissue and point to biology that is accessible only with spatial measurement. Another less explored source of spatial variation can arise from the assay itself. Spatial platforms that use imaging-based techniques vary in their ability to detect transcripts above a putative noise level, which has a direct consequence on the observed spatial patterns (Plummer et al., 2026). Additionally, assays that rely on microarray-based capture are known to be susceptible to positional differences in RNA capture efficiency that can induce correlation in observed counts (Mazoure et al., 2017; Normandeau et al., 2021). When unaccounted for, this structure is vulnerable to being conflated with biological processes in downstream analysis. In light of this, it can be difficult to determine whether observing a spatial pattern means that the spatial context is necessary to explain it.

Existing methods for SVG gene identification largely attribute spatial terms to an unspecified process rather than to a specific generative mechanism (Yan et al., 2025). Common approaches fit a Gaussian process to normalized counts to identify spatial covariance (Svensson et al., 2018; Weber et al., 2023). Other methods incorporate count distributions into generalized linear mixed models, but spatial relationships are modeled as unconstrained nuisance terms that inform hypothesis tests or differential expression (Ospina et al., 2024; Sun et al., 2020). A more recent approach aims to separate technical from biological spatial variation (Taychameekiatchai et al., 2026), but does so by using assigning labels to individual genes that are difficult to verify without information orthogonal to the transcriptomic measurements. Thus, the presence of a spatial term in a model does not on its own reveal whether a detected pattern reflects known domain structure, a shared technical process, or gene specific biology. These distinctions require additional information about the generative process that connects plausible biological and technical processes.

To address these limitations, we present a framework that unifies sources of biological and technical variation. Our method, called stipple, explicitly models count data as a bursty, stochastic biological process followed by spatially smooth capture. The goal is to discriminate between genes whose expression pattern is fully explained by domain or by a spatial technical process, and those whose expression that suggests context-dependent biology that persists beyond known tissue organization. Our proposed framework adapts mechanistically motivated observation models of count data to allow for a correlated molecule capture process. The estimated gene parameters can be interpreted in terms of their relationship to the transcriptional mechanism described in the model, while the estimated capture field can be used to condition residual expression after domain specific effects have been removed. We show that DVGs can be identified by interrogating the gene likelihood parameters and SVGs can be classified by assessing spatial autocorrelation before and after conditioning on the estimated capture field. Overall, this approach provides a starting point for separating sources of spatial variation and incorporating more complex transcriptional and measurement processes to inform inference in various spatial contexts.

## Results

### stipple infers technical and biological parameters in spatial transcriptomics data

A SRT dataset consists of a mRNA count matrix together with a spatial index for each unit of observation. There are several strategies for encoding spatial coordinates (Moses and Pachter, 2022), though the Visium spatial platform from 10X Genomics has seen wide adoption. In this assay, capture probes containing a positional barcode are immobilized on a functional surface (Fig. 1A). After tissue permeabilization, transcripts diffuse from the tissue and are captured by nearby probes, which allows transcripts to be assigned back to their approximate physical location after sequencing. When tissue annotations are available, the stipple model learns the capture rate at each location together with gene-likelihood parameters that describe the transcriptional process (Fig. 1A). The output of the model is the inferred technical capture field along with parameter matrices corresponding to the learned mean and dispersion for each gene and domain (Fig. 1B).

**Fig. 1.**
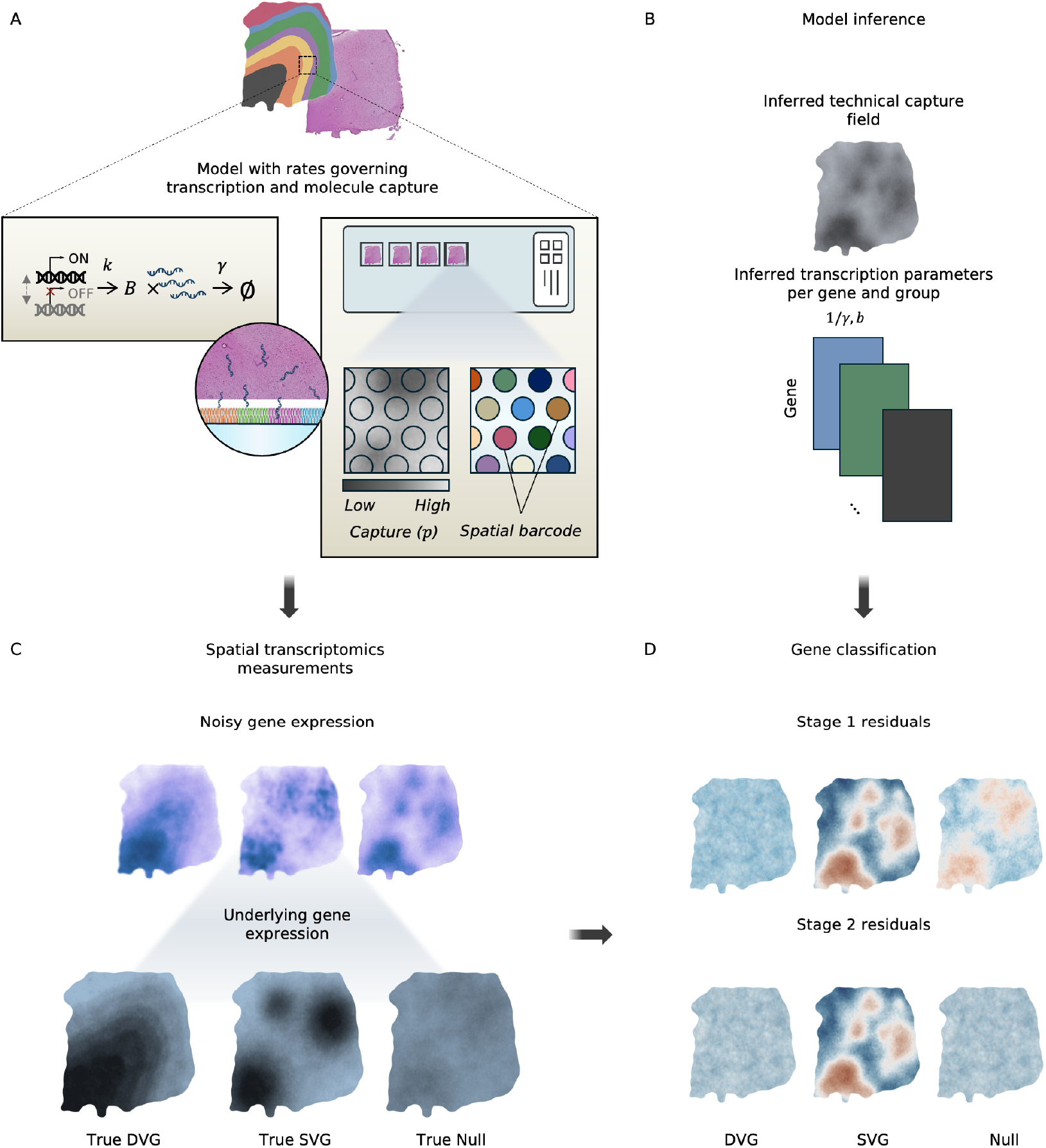
Overview of the stipple model. A.) For simplicity, the example considers a spatial transcriptomics dataset measured using the 10X Genomics Visium assay. The stipple model assumes that each location in the assay has a position-dependent capture rate and that each observation within a group has the same transcriptional rate parameters. Counts are modeled using a bursty model of transcription, which induces a Negative Binomial distribution at steady state. B.) The model outputs a prediction of the technical spatial field and the distributional parameters. C.) The observed counts are a combination of the biological and capture process. D.) The estimated field can be used to classify genes based on their expression relative to the estimated field and group annotations.

Following work in single cell RNA-seq (scRNA-seq) analysis, we model transcript abundance using the well characterized bursty model of transcription (Singh and Bokes, 2012; Gorin and Pachter, 2022; Gorin et al., 2022). At steady-state, this process gives rise to a Negative Binomial distribution parameterized by shape *k/γ* and scale *b*, where *k* and *γ* are the rates of transcription and degradation, respectively, and *b* is the burst size (see Methods). Under Bernoulli a sampling noise model, the observed distribution of counts at each location remains Negative Binomial, except that the burst size is scaled by the Bernoulli parameter *p*. The scale of the average capture rate is anchored beforehand so that each location’s technical capture rate is identifiable.

To represent structured technical noise, we allow *p* to vary smoothly across the observed capture area (Fig. 1A-B). Under this model, spatial correlation in *p* can induce autocorrelation in the observed counts, even when the underlying distributions in each domain have similar underlying biology. However, each dataset provides only one observation of the latent spatial field. To address this, we constrain the covariance structure of the capture field so that information is shared between observations for inference. Consequently, spatially patterned gene expression can arise because of differences in biological parameters, as is the case for DVGs, because of correlation with the technical field, or from a combination of these mechanisms. Gene specific spatial variation represents a case where neither source explains the remaining variation.

The gene classification provided by stipple guides exploratory biological analysis by linking different spatial patterns to different questions (Fig. 1C). DVGs can be explored within the regions that they identify, whereas residual SVGs prioritize genes for exploring expression gradients or previously unannotated spatial organization. Genes whose spatial patterns are explained by the shared capture field can instead be deprioritized as candidates for spatially dependent or gene-specific biology (Fig. 1D).

### A spatially-resolved capture field improves SVG discrimination in simulated data

To assess the ability of the stipple model to recover a known spatial field and ground truth gene class labels, we simulated a transcriptomics dataset with positional capture bias and known group annotations (Fig. 2A-B). Briefly, gene counts for cell *i* in group *k*(*i*) were broadly divided into categories including, (1) null genes with stable expression across groups, (2) DVGs with differences in mean expression level between groups, (3) genes with spatial variation associated with the shared spatial field, and (4) genes with spatial variation independent of the shared field (Fig. 2D). After subtracting the group-specific effects from the counts, we asked whether our approach could correctly identify when the residual spatial pattern was aligned with the technical bias. Genes were called SVGs using a threshold of Moran’s *I >* 0.2.

**Fig. 2.**
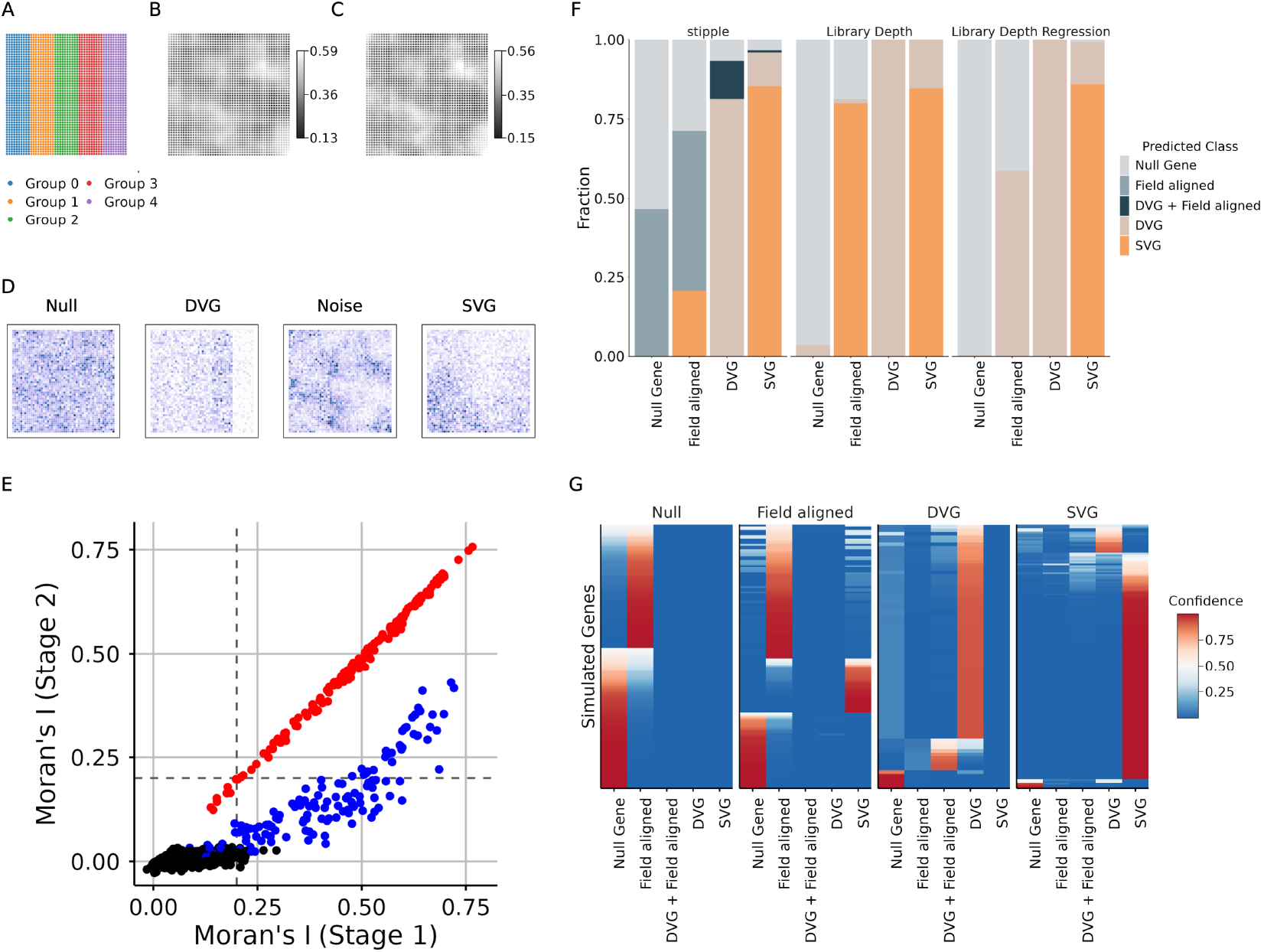
We devised a simple simulation consisting of A.) 5 ‘biological’ groups observed on a B.) spatially-biased capture field. C.) The model predicted the underlying capture field. D.) Different classes of genes were simulated from the model and subsequently classified. E.) The two-stage process, which includes conditioning the observed gene residuals on the estimated field. Genes that maintain spatial autocorrelation after conditioning are predicted to be SVGs. F.) We compared the predicted gene class to the true gene class. Only the stipple model can distinguish SVGs from field-aligned genes. G.) The confidence of the class predictions in panel E. generated from the stipple model. Each panel is labeled on the top by the true gene class assignment.

First, stipple recovered the expected shape of the simulated technical field (Fig. 2B-C). By construction, group differences in mean capture efficiency are absorbed into the gene-specific expression estimates. In simulations, differences in average capture efficiency between groups were small, so the number of genes called DVGs based on the fitted *µ*_*gk*_ was not inflated in the simulation analysis (Fig. 2E-F). In real data, where group differences in capture rate are unknown, the compositional mean 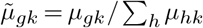 can be used as a diagnostic for group specific scaling effects when calling DVGs. Since 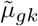 is invariant to multiplicative shifts in the capture rate, differences in 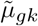 represent gene specific changes in relative abundance. Importantly, the stipple model rarely assigned field-aligned or null genes to gene categories that would suggest further analysis (i.e. DVG or SVG) (Fig. 2F). We assigned a confidence score to each classification call to assess whether the mislabeled gene classes were mostly borderline or hard to call genes (Fig. 2G). Perhaps unsurprisingly, the field-aligned and null genes showed considerable overlap. One explanation for this is that the variance of low expression null genes is dominated by the technical field component. In any case, the broader assignment for these genes as being neither DVG nor SVG is consistent with the simulation.

Typical approaches for identifying candidate SVGs do not target gene classification as an inferential goal. Instead these methods focus on the binary task of calling genes as SVG or not. Therefore, we asked whether common preprocessing workflows are sufficient prevent field-aligned genes from being flagged as candidate SVGs. We compared our estimation framework to library size normalization and to regressing the log1p transformed counts on the log-transformed library size. Though both of these approaches should ‘normalize out’ global scaling factors, both of these approaches classify most field-aligned genes as SVGs or DVGs (Fig. 2F). In contrast, regressing each gene’s group mean-centered Pearson residual on the estimated field selectively reduces the Moran’s *I* of field-aligned genes, while leaving the Moran’s *I* of field independent genes relatively unchanged. Thus, we conclude that this workflow effectively separates DVGs from SVGs, and can accurately identify genes whose spatial pattern is biased by a global technical scaling factor.

### DLPFC Visium data support a shared, spatially structured scaling field

Next, we set out to determine whether the model fitting procedure we described could be used to facilitate accurate SVG detection and DVG classification in real SRT data. We applied the stipple model to a human dorsolateral prefrontal cortex (DLPFC) dataset generated on the 10X Genomics Visium V2 platform (Maynard et al., 2021). Briefly, 12 samples from 3 individuals were assayed across 3 Visium spatial arrays. For each individual, two anterior DLPFC samples and two posterior DLPFC samples were assayed on separate capture areas on the same array. Since a true reference for the capture field is not available, this experimental design provides a useful setting for evaluating technical variation because adjacent tissue sections are expected to share broad anatomical organization and properties such as average cell size within corresponding tissue compartments.

### Assessing alternate interpretations of the inferred spatial field

Before assessing the results, we explored the correlation between the estimated field and several plausible biological interpretations. The stipple model parameterizes the field as a spatially coordinated capture process that is technical in origin. Since no spatially resolved external RNA standard was available for these data, the biological and technical contributions to global scaling cannot be identified from the observed counts alone. Consequently, we examined whether the properties of the inferred field could be consistent with our technical parameterization.

We fit the model to each sample separately and found support for a global scaling factor across samples. As in the simulated data, the estimated field was robust to the composition of genes used for model fitting (Supp. Fig. S1). Similar spatial structure was recovered after excluding genes contributing the largest fractions of observed transcripts, indicating that estimation of the field was not driven by a small number of highly abundant genes (Supp. Fig. S1).

Using sample 151673 as a representative example (Fig. 3A), we observed that while the total counts and estimated field are correlated, their spatial patterns are distinct (Fig. 3B-C). Specifically, we observed that locations with relatively high total counts did not consistently overlap with regions of high capture (Supp. Fig. S2). This behavior can be expected if the total counts observed at each location are a measure of an underlying biological process in addition to variable capture across the assay.

**Fig. 3.**
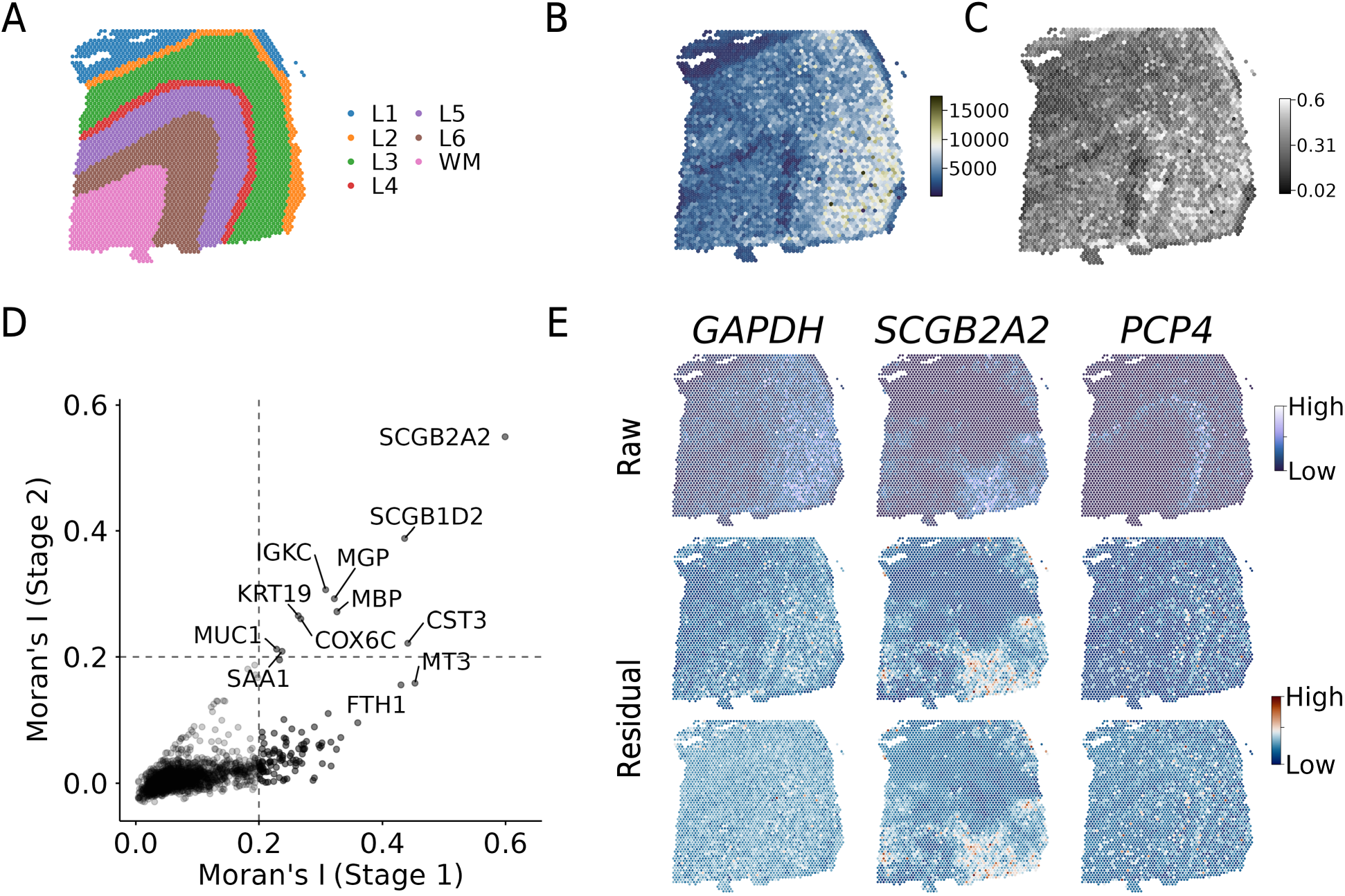
A. Annotated layers in sample 151673. B-C.) The total counts and estimated spatial field have distinct spatial patterns. D. Moran’s I differentiates gene classes after regressing the Pearson residual on the estimated field. Field-independent SVGs retain their spatial autocorrelation in the second stage. E. *GAPDH* is an example of a field-aligned SVG that loses spatial correlation in stage 2. On the other hand, *SCGB2A2* retains spatial autocorrelation in both stages. *PCP4* is an example of DVG identified via DE analysis with the estimated group mean parameter.

Another possibility is that the estimated field is a proxy for cell size or RNA abundance. We compared the estimated field to the fluorescent cDNA images provided in the supplement from the original authors (Supp Fig. S8; Ref (Maynard et al., 2021)). Interestingly, the cDNA in tissue slices from two samples (Br5595, Br8100) appeared homogeneous throughout the tissue, while sample from Br5292 showed reduced cDNA in L1 and white matter layers, which is consistent with known biology for this region of the brain (Bruno et al., 2022). Upon visual inspection, there was not a strong correspondence between the cDNA staining and the predicted field. In samples Br5595 and Br8100, the spatial field exhibited considerable spatial variability in regions on relatively constant cDNA abundance. This observation lessens support for a cell size interpretation of the spatial field. We explored this further using a metric for cell density. Since the Visium platform does not have single cell resolution, spots with containing more than one cell are expected to have a greater local abundance of RNA. We observed that the inferred field did not correlate strongly with the number of nuclei overlapping each spot, which we used as a proxy for cell density (Supp. Fig. S2). While this is not a direct measure of cell size or total RNA per spot, this result argues against a simple mechanism where differences in cellular abundance reproduce the field. Next, we asked whether converting the counts to relative abundance led to a reduction in the correlation between the expression and estimated field, since any shared multiplicative factor cancels in the fraction. Indeed, there was observed broad attenuation of the correlation between gene expression and the estimated field when we converted the counts to relative abundance (Supp. Figs. S4-S5). The effect was stronger in genes that were strongly aligned with the spatial field. This is consistent with a global scaling factor rather than a coordinated biological process that affects only a subset of genes.

In the absence of orthogonal measurements, we can not exclude biological scaling. However, a biological mechanism would need to operate continuously within domains, remain largely unexplained by cell density, broadly affect unrelated gene classes, and alter total transcript count without producing corresponding gene-specific expression programs. We conclude that the inferred field is at least in part consistent with spatially heterogeneous technical capture efficiency.

### Explicit modeling separates classes of expression spatial variation

In the DLPFC, differences in mean expression across the cortical layers for several genes provide a well-characterized axis of spatial variation, and these group-level expression differences alone are sufficient to produce spatially variable expression (Maynard et al., 2021). As exploratory analysis before fitting the model, we computed the Pearson residual from the observed group mean within each annotated layer. Across samples, many genes retained positive spatial autocorrelation following group-mean centering, and the spatial pattern of the residuals was qualitatively similar. Indeed, singular value decomposition of the residuals revealed a dominant rank-1 component whose loading scaled with each gene’s mean (Supp. Fig. S3). Thus, the transcriptional differences between cortical layers do not fully account for the spatial variation of expression in these data. Instead, spatial variation persists after the dominant discrete anatomical structure is removed.

To distinguish DVGs from genes whose residual spatial structure reflects the estimated shared field and from genes with unexplained spatial variation, we computed the observed Pearson residuals using the observed within-group mean and variance for each gene (Fig. 3E). We conditioned these residuals on the estimated field and used the regression residual to re-compute Moran’s *I* (Methods). First, we observed that DVGs are not generally called SVG at the first stage, since the residual accounts for domain-level effects (Supp. Fig. S7). Instead, since stipple recovers absolute expression rather than a normalized quantity, DVGs can be recovered by computing the log fold-change between the estimated gene-level and layer-level parameters. Again using sample 151673 as an example, the fraction of genes called SVGs reduced after the residual correction. Genes whose apparent SVG status was attributable to field included widely expressed ribosomal genes and genes such as *GAPDH* which are associated with homeostatic cellular processes (Fig3E) (Sirover, 2005). Genes retaining independent spatial structure included *SCGB2A1* and *SCGB2A2*, normally associated with glandular and secretory tissue (Jackson et al., 2011) and plausibly reflect expression that is not well explained by the typical DLPFC organization (Fig. 3E). The DVG category recovers known layer marker genes, such as *PCP4* in Layer 4/5 (Fig3H) (Renelt et al., 2014).

We compared stipple-called SVGs to those identified by SpatialDE (Svensson et al., 2018). The methods vary in their agreement on the genes that are neither SVGs nor DVGs (Supp. Fig. S6), though it is possible that the log1p transformation in the SpatialDE workflow induces spurious spatial gradients and differences between groups (Lun, 2018; Booeshaghi et al., 2022). SpatialDE called most DVGs as spatially variable, which can be expected as the processing workflow for SpatialDE neither removes genespecific nor group-specific effects from the counts (Svensson et al., 2018). As in the simulated results comparing normalization strategies, genes that have a combination of field-associated spatial patterns are typically called SVGs, whether or not there is an additional group-level or genelevel effect (Supp. Fig. S6). Together, these results show that the value of explicitly modeling spatial scaling is the ability to preserve biological abundance while distinguishing spatial variation that is shared across the transcriptome from spatial structure specific to individual genes.

## Discussion

The stipple model jointly estimates a latent, spatially structured technical capture field and group-level gene expression parameters from SRT count data. The observation that spatial variation in gene expression can arise from biological and technical processes requires a framework to explicitly model both sources of variation. Thus, we investigated whether the spatial signal in gene expression counts could be attributed to a shared, technical source. Our simulations showed that without explicit modeling of the shared source of spatial variation, genes whose residual spatial pattern is a consequence of the technical field cannot be readily distinguished. Explicitly fitting the shared capture field resolved this ambiguity when compared to standard data processing workflows. In both the simulated and DLPFC data, several genes initially called spatially variable can be understood to have variability driven by group-level differences and field-level artifacts, and only a small number of genes were found to retain their independent spatial structure.

A central limitation of our work is that gene expression counts alone cannot uniquely decompose the shared field into technical and biological components. Specifically, a transcriptome-wide multiplicative field could arise from biological differences in cell size, total RNA content, or global transcriptional activity that alters the abundance of many transcripts simultaneously. Distinguishing these mechanisms experimentally is challenging in part because many SRT assays are destructive and true technical replicates of the same tissue slice are unavailable. Adjacent sections provide useful information and can be considered “pseudoreplicates”, but differences in sectioning depth, cellular cross-sections, tissue alignment, and section processing can all result in non-trivial differences between even closely spaced sections. Spatially-resolved RNA standards are an attractive solution for more direct validation, but their use would require careful consideration of a delivery system with a known spatial distribution across the capture area.

In the absence of a direct measurement of capture efficiency, we evaluated the estimated field against properties expected under various biological explanations. We found that the estimated field is robust to the gene subset use for model fitting, exhibited spatial structure distinct from total counts, was not explained by a proxy for cell density, demonstrated broad attenuation of gene association after conversion to relative abundance, and aligned with a spatial axis orthogonal to the annotated organization. Each of these observations narrows the biological mechanisms that can produce a spatially patterned, transcriptome-wide scaling within tissue domains. A spatial gradient in cell size remains an alternative and cannot be excluded with present data. Direct comparison of the inferred field with a spatially resolved external standard will be important for establishing the relative technical and biological contributions to this shared spatial variation.

Furthermore, the choice to require spatial annotations is another consequence of the observation that group-level expression parameters and location-level capture estimates are not uniquely identifiable. Typical analysis workflows identify SVGs and use them to define tissue domains, even though a spatially structured technical process affects both the scale and apparent spatial organization of the observed counts. Importantly, including annotations in the fitting process enables identification of a spatial variation that is attributable to tissue organization a priori. However, the model cannot detect cases in which two annotated groups share similar mean expression for a gene but differ in absolute expression in a way that is compensated by differences in local capture efficiency. Still, the ability to model tissue organization directly is particularly relevant when the analysis goal is inference of spatial regulation within an already characterized tissue. Indeed, once broad domains are known, learning them through SVG analysis provides little new information. In contrast, identifying continuous spatial variation that does not respect domain boundaries or operates within a distinct compartment can reveal spatial variation patterns that are otherwise obscured by the programs that distinguish tissue domains. The stipple model therefore treats annotations as an important biological prior that allows spatial information to be directed towards inference of variation that remains unexplained by known anatomy.

More broadly, our results suggest that normalization and spatial inference should not be separated into distinct analysis problems. Although normalizing counts with library size can correct the technical scaling factor, it obscures the absolute differences between tissue domains, removing valuable biological information. Furthermore, ignoring the shared field can result in confounding it with gene-specific biology. Explicitly decomposing sources of spatial variation in a unified model provides a framework toward assigning spatial expression patterns to specific biological and technical mechanisms.

## Acknowledgments

Thanks to members of the Pachter lab who provided valuable input in discussions related to the methods that led to the results in this paper. KJ was partially funded by the Caltech Cell Interaction Initiative.

## Author contributions

KJ implemented the method, generated the results, and drafted the manuscript. GG conceived of the structure of the capture model and derived the observation model for the counts. LP helped develop the method. All authors reviewed and edited the manuscript.

## Methods

### Motivation

Spatial transcriptomics data consist of gene expression counts measured at discrete tissue locations. At each location *i* = 1,…, *n*, counts for gene *g* = 1,…, *G* are observed, and the spatial differences in the observed counts may reflect both biological signal and technical processes. Biological signal can arise from several mechanisms. In one case, cell-cell communication and coordination of gene programs can produce global and/or local continuous gradients of expression across a tissue. This signal can also reflect the preference of certain cell types to migrate to distinct regions. In the latter case, spatial differences are largely attributable to differences between discrete biological groups and can, in principle, be detected without explicitly modeling spatial relationships among locations.

A less explored mechanism for spatial variation in SRT data arises from technical heterogeneity in capture efficiency across the spatial capture assay. This mechanism can both obscure true spatial variation and produce artificial spatial correlations that can be misinterpreted as biological signal. Here, we explicitly model capture efficiency as a spatially varying technical process while allowing gene expression parameters to differ between annotated biological groups. This decomposition separates expression differences attributable to known discrete biological structure from spatial variation shared across genes through the capture process. Consequently, after accounting for annotated group differences and spatially structured capture efficiency, residual spatial expression patterns provide evidence for biological variation that cannot be explained by discrete tissue annotations or by the modeled technical capture process.

### Bursty Transcription and Bernoulli Capture Model

We model transcription of a single molecular species using a bursty model of transcription. The bursty model can be derived using the Chemical Master Equation (CME) formalism, which describes the time evolution of the probability distribution over discrete molecule counts (Singh and Bokes, 2012; Gorin et al., 2022).

Let *X*_*g*_ denote the steady-state count of mRNA molecules for gene *g*. We assume that molecules are produced in bursts at rate *k*_*g*_, where each burst produces a geometrically distributed number of molecules with mean *b*_*g*_. Molecules are degraded at rate *γ*_*g*_. It has been shown that this bursty production and degradation process results in a Negative Binomial steady-state distribution (Singh and Bokes, 2012).

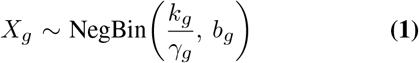

with mean *µ*_*g*_ = *k*_*g*_*b*_*g*_*/γ*_*g*_ and variance 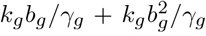. At steady state, the transcription rate *k*_*g*_ and degradation rate *γ*_*g*_ are not independently identifiable from the marginal count distribution. We therefore define the degradation rate relative to transcription rate, *γ*_*g*_*/k*_*g*_. Without loss of generality we set *k*_*g*_ = 1, so that *ϕ*_*g*_ = 1*/γ*_*g*_ and the steady-state distribution is

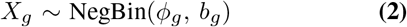

with mean *µ*_*g*_ = *ϕ*_*g*_*b*_*g*_ and variance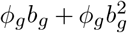. In the case where location *i* is assigned to group *k*(*i*) according to discrete labels provided with the data, the expression of gene *g* can vary by group-specific gene parameters. This amounts to adding an index *k* to parameters *ϕ*_*g*_ and *b*_*g*_, which we have omitted for simplicity.

#### Capture Model

Observed counts reflect molecules that are successfully captured from the true transcript population. We model capture as a Bernoulli capture process in which each molecule at location *i* is independently captured with probability *p*_*i*_ *∈* (0, 1]. Under Bernoulli capture, the resulting distribution for observed counts *Y*_*ig*_ at location *i* and gene *g* is

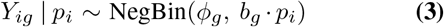

with mean E[*Y*_*ig*_ |*p*_*i*_] = *ϕ*_*g*_*b*_*g*_*p*_*i*_ and variance Var(*Y*_*ig*_| *p*_*i*_) = *ϕ*_*g*_*b*_*g*_*p*_*i*_ + *ϕ*_*g*_(*b*_*g*_*p*_*i*_)2. Bernoulli capture of a Negative Binomial random variable scales only the burst size *b*_*g*_ by the capture probability *p*_*i*_.

Equivalently, writing *µ*_*g*_ = *ϕ*_*g*_*b*_*g*_ for the pre-capture mean expression, the observation model has mean E[*Y*_*ig*_ | *p*_*i*_] = *µ*_*g*_ *p*_*i*_ and variance *µ*_*g*_*p*_*i*_ + (*µ*_*g*_*p*_*i*_)^2^*/ϕ*_*g*_, where *ϕ*_*g*_ is the dispersion. This mean–dispersion (*µ*_*g*_, *ϕ*_*g*_) parameterization, with a group index (*µ*_*g,k*_, *ϕ*_*g,k*_) when group-level parameters are estimated, is the form used in the inference model (see Methods, section ‘Model Inference’) and in the Pearson residual defined below.

A Poisson likelihood is available as a special case in the limit *ϕ*_*g*_ *→ ∞* with *b*_*g*_ *→* 0 such that *ϕ*_*g*_*b*_*g*_ = *λ*_*g*_ remains fixed, giving *Y*_*ig*_ | *p*_*i*_ ~ Pois(*λ*_*g*_*p*_*i*_).

### Spatial Capture Field

To explain the observed spatial correlation in the count data, capture probabilities *p*_*i*_ are modeled as spatially correlated across locations, and the capture probability is shared across genes at a given location. This spatial correlation is induced by representing the underlying latent field as a weighted combination of a fixed set of compactly-supported basis functions centered at spatial knots, so that nearby locations that share overlapping basis support are constrained to have similar values.

#### Knot Placement

A target number of knots is user-specified, subject to a lower bound of *M/n ≥* 0.5 that ensures the basis is sufficiently dense relative to the number of observed locations to represent meaningful spatial variation in capture rate. Knots are placed on a regular hexagonal grid over the bounding box of the tissue spatial coordinates, with the knot spacing set to realize the target number of knots. Fields with structure at fine spatial scales relative to the sampling resolution may require a substantially larger *M/n*.

#### Basis Function Construction

Each knot *m* has an associated compact-support radius *r*_*m*_, set to the knot spacing scaled by an overlap factor of 1.5. The raw basis matrix **B**^raw^ *∈* ℝ^*n×M*^ is constructed using a bisquare radial kernel function:

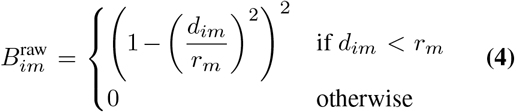

where *d*_*im*_ = ∥ coords_*i*_ *−* knot_*m*_∥ is the Euclidean distance between location *i* and knot *m* in data coordinates. The compact support of the bisquare kernel makes **B**^raw^ sparse, so that each location receives nonzero contributions only from knots within radius *r*_*m*_. Each row of **B**^raw^ is normalized to sum to one, yielding a partition of unity basis matrix **B** *∈* ℝ^*n×M*^ :

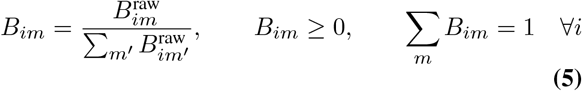

#### Latent Field and Capture Rate

A weight vector **w** *∈* ℝ^*M*^ is assigned to the knots. The latent spatial field **u** = (*u*_1_, …, *u*_*n*_)^⊤^ at each location is the basis-weighted sum:

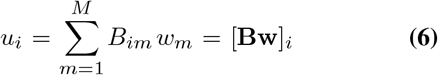

The partition of unity property (Equation 5) ensures *u*_*i*_ is a convex combination of the weights, so the spatial variation in *u*_*i*_ is controlled entirely by the spread of **w** across knots.

#### Group-level centering

When group-level parameters are estimated, each location’s expected count is *λ*_*ig*_ = *µ*_*g,k*(*i*)_ *· p*_*i*_, which is a product of a group-level expression and a location-level capture rate. Here, a genuine between-group difference in total expression and a spurious between-group difference in average capture efficiency are both consistent with the same observed counts. When group membership correlates with spatial position, as is typical for spatially clustered tissue domains, the latent field weights **w** can absorb systematic between-group offsets. To remove this confound structurally, the latent field is centered within each group on the logit scale before conversion to a capture probability. Letting *Z ∈ {*0, 1*}* ^*n×K*^ denote the one-hot-encoded group indicator matrix, the group-mean centered field is:

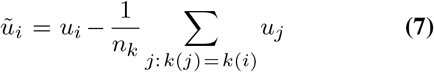

where *n*_*k*_ = Σ_*j*_ I[*k*(*j*) = *k*] is the number of locations in group *k*. Since centering commutes with the linear map **Bw**, this is algebraically equivalent to replacing **B** with the group-mean-centered basis:

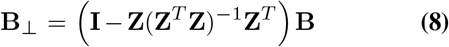

Because the columns of **Z** are mutually orthogonal, **Z**^*T*^ **Z** is diagonal and the projection reduces to per-group mean centering of each basis column. The centering is applied to *u*_*i*_ after forming **Bw** rather than to **B** itself, which preserves the exact sparsity of **B**.

Finally, the capture probability at location *i* is defined as

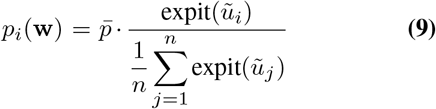

where 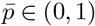 is a user-supplied known average capture rate and expit(*x*) = (1 + *e*^*−x*^)^*−*1^. The normalization by the empirical mean of the latent field enforces 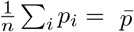 exactly for any value of **w**. When **w** = **0**, expit(*u*_*i*_) = 0.5 everywhere and 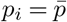 at every location, so the natural origin of the weight prior corresponds exactly to a spatially uniform capture rate equal to the known mean.

#### Estimation of Average Capture Probability

The absolute scale of the capture probability is not identifiable from the count likelihood alone because the expected count depends on the product of the biological expression level and the location-specific capture probability, *E*[*Y*_*ig*_ | *p*_*i*_, *k*_*i*_] = *µ*_*gk*_*i p*_*i*_. Consequently, proportional changes in *p*_*i*_ can be offset by proportional changes in *µ*_*gk*_ without changing the expected counts. We therefore use the observed total counts to obtain a relative estimate of capture efficiency and impose a probability constraint to calibrate its absolute scale.

Let *T*_*i*_ = Σ_*g*_ *Y*_*ig*_ denote the total observed transcript count at location *i*. Under the multiplicative capture model,

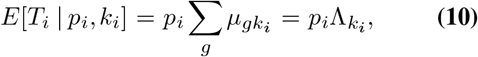

where Λ_*k*_ = Σ_*g*_ *µ*_*gk*_ denotes the underlying total transcript abundance for biological group *k*. Therefore, variation in *T*_*i*_ reflects both variation in capture probability and differences in total biological transcript abundance among groups.

Let *n*_*k*_ denote the number of locations assigned to group *k, N* = Σ_*k*_ *n*_*k*_, and 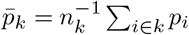. The expected mean total count across all locations is

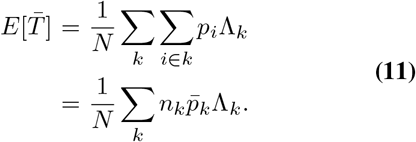

We assume that the average capture probability is common across biological groups, such that 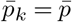 for all *k*. Under this assumption,

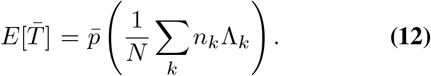

Define the observed library-size ratio 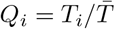. Its expected value is approximately

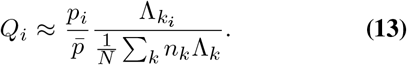

Thus, *Q*_*i*_ alone does not estimate relative capture efficiency when the biological total-expression scale Λ_*k*_ differs among groups.

To remove this group-level biological contribution, let 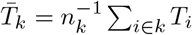 denote the observed mean total count within group *k*. Under the common mean-capture assumption,

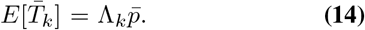

Consequently, the ratio of a location’s total count to the mean total count of its biological group provides an estimate of capture efficiency relative to the common mean,

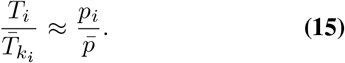

Equivalently,

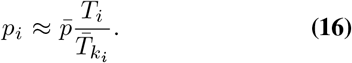

The observed counts identify this relative spatial scale but do not identify the absolute value of 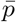. Because *p*_*i*_ represents a probability, it must satisfy 0 *< p*_*i*_ *≤* 1. Therefore,

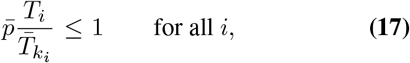

which implies

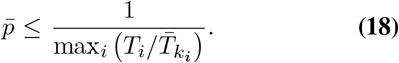

We calibrate the absolute scale by selecting the largest value of 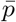 consistent with this constraint. The estimated average capture probability is therefore

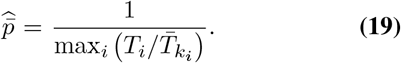

Importantly, the constraint max_*i*_ *p*_*i*_ = 1 is a calibration assumption rather than a quantity identified from the observed counts. In particular, the data do not establish that any location captures all RNA molecules available for measurement, and it is likely the case that max_*i*_ *p*_*i*_ *<* 1. Thus, the estimated 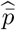 defines a maximum absolute scale of the latent capture field under this reference. Importantly, the shape of the estimated field within each group is recoverable no matter the choice for this calibration. Also, because total RNA abundance may vary across annotated groups, the total count ratios are not unbiased, so their use is restricted to providing a heuristic anchor for the otherwise unidentified scale.

### Model Inference

#### Bayesian Model

The model is specified as a joint distribution over observed counts and all parameters, enabling maximum a posteriori (MAP) estimation by maximizing the penalized log-joint. The full model is defined as

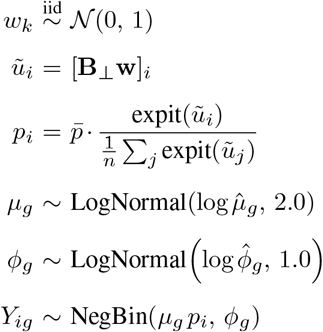

The parameters 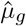 and 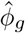 are plug-in estimates computed once from the data before fitting, and **B**_*⊥*_ is the groupmean-centered basis matrix (Equation 7). The hyperpriors on *µ*_*g*_ and *ϕ*_*g*_ are weakly informative on the log scale and permit several orders of magnitude of deviation from the plug-in center.

#### Optimization

Parameters are estimated by MAP optimization of the penalized log-joint likelihood. The parameters to be estimated are the spatial basis coefficients *w*, the gene-specific pre-capture means *µ*_*g*_, and, for the Negative Binomial likelihood, the gene-specific dispersion parameters *ϕ*_*g*_. Parameters *µ*_*g*_ and *ϕ*_*g*_ are optimized in log-space to enforce positivity constraints.

The negative penalized log-joint is minimized using the Adam optimizer (Kingma and Ba, 2017) with gene minibatches of size *G*^*′*^ = 200 genes per iteration. All *n* locations contribute to every gradient step. Optimization proceeds for up to 3, 000 iterations and terminates early if the relative change in the objective over a rolling window of 20 consecutive iterations falls below 10^*−*6^.

### Classification of Spatially Variable Genes

After fitting, spatially variable genes are identified by conditioning the observed Pearson residuals on the estimated spatial field from the model. For gene *g* at location *i* the Pearson residual is

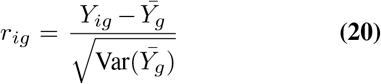

computed separately within each annotated group. A gene’s residual vector **r**_*g*_ may retain spatial autocorrelation attributable to the shared capture field even after group-mean centering, since the field is common across genes and its residual imprint scales with each gene’s fitted mean. To distinguish field-attributable residual structure from spatial variation independent of the field, we regress **r**_*g*_ on the fitted capture rate 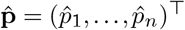 and take the regression residual 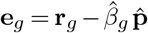, where 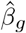 is the ordinary least squares coefficient. This regression tests whether a gene’s residual spatial pattern is a rescaled copy of the fitted field. If so, **e**_*g*_ is spatially flat, while genuine field-independent spatial structure survives in **e**_*g*_.

Spatial autocorrelation of both **r**_*g*_ (Stage 1) and **e**_*g*_ (Stage 2) is quantified using Moran’s *I* (Moran, 1950), with a row-standardized symmetric *k*-nearest-neighbor spatial weight matrix (*k* = 6). Since *n* is large, even slight spatial auto-correlation produces a significant *p*-value under a permutation or normal-approximation null. Thus we base a gene’s spatial-variability call on the magnitude of *I* rather than on its significance. A gene is called spatially variable if *I*(**e**_*g*_) *>* 0.2. The same statistic computed on **r**_*g*_ prior to field regression is reported separately to quantify the fraction of apparent spatial variability attributable to the shared field.

## Data availability statement

The DLPFC Visium data are available from the original authors in the paper cited in this manuscript (Maynard et al., 2021). We accessed the raw data at the following link: https://github.com/LieberInstitute/HumanPilot.

## Code availability

The code implementing the stipple workflow is available as a pip installable package at https://github.com/pachterlab/stipple/. Code and data for reproducing manuscript results can be found at https://github.com/pachterlab/JGP_2026/.

## References

A. Sina Booeshaghi, Ingileif B. Hallgrímsdóttir, Ángel Gálvez-Merchán, and Lior Pachter. Normalization for sampled count data, May 2022.

Ariane Bruno, Sebastian Bludau, Hartmut Mohlberg, and Katrin Amunts. Cytoarchitecture, intersubject variability, and 3D mapping of four new areas of the human anterior prefrontal cortex. Front. Neuroanat., 16:915877, August 2022. ISSN 1662-5129. doi: 10.3389/fnana.2022.915877.

Gennady Gorin and Lior Pachter. Modeling bursty transcription and splicing with the chemical master equation. Biophysical Journal, 121(6):1056–1069, March 2022. ISSN 0006-3495. doi: 10.1016/j.bpj.2022.02.004.

Gennady Gorin, John J. Vastola, Meichen Fang, and Lior Pachter. Interpretable and tractable models of transcriptional noise for the rational design of single-molecule quantification experiments. Nature Communications, 13(1):7620, December 2022. ISSN 2041-1723. doi: 10.1038/s41467-022-34857-7.

Brian C. Jackson, David C. Thompson, Mathew W. Wright, Monica McAndrews, Alfred Bernard, Daniel W. Nebert, and Vasilis Vasiliou. Update of the human secretoglobin (SCGB) gene superfamily and an example of ‘evolutionary bloom’ of androgen-binding protein genes within the mouse Scgb gene superfamily. Human Genomics, 5(6):691, October 2011. ISSN 1479-7364. doi: 10.1186/1479-7364-5-6-691.

Diederik P. Kingma and Jimmy Ba. Adam: A Method for Stochastic Optimization, January 2017.

Aaron Lun. Overcoming systematic errors caused by log-transformation of normalized singlecell RNA sequencing data, August 2018.

Kristen R. Maynard, Leonardo Collado-Torres, Lukas M. Weber, Cedric Uytingco, Brianna K. Barry, Stephen R. Williams, Joseph L. Catallini, Matthew N. Tran, Zachary Besich, Madhavi Tippani, Jennifer Chew, Yifeng Yin, Joel E. Kleinman, Thomas M. Hyde, Nikhil Rao, Stephanie C. Hicks, Keri Martinowich, and Andrew E. Jaffe. Transcriptome-scale spatial gene expression in the human dorsolateral prefrontal cortex. Nature Neuroscience, 24(3): 425–436, March 2021. ISSN 1546-1726. doi: 10.1038/s41593-020-00787-0.

Bogdan Mazoure, Robert Nadon, and Vladimir Makarenkov. Identification and correction of spatial bias are essential for obtaining quality data in high-throughput screening technologies. Sci Rep, 7(1):11921, September 2017. ISSN 2045-2322. doi: 10.1038/s41598-017-11940-4.

Christopher R. Merritt, Giang T. Ong, Sarah E. Church, Kristi Barker, Patrick Danaher, Gary Geiss, Margaret Hoang, Jaemyeong Jung, Yan Liang, Jill McKay-Fleisch, Karen Nguyen, Zach Norgaard, Kristina Sorg, Isaac Sprague, Charles Warren, Sarah Warren, Philippa J. Webster, Zoey Zhou, Daniel R. Zollinger, Dwayne L. Dunaway, Gordon B. Mills, and Joseph M. Beechem. Multiplex digital spatial profiling of proteins and RNA in fixed tissue. Nat Biotechnol, 38(5):586–599, May 2020. ISSN 1546-1696. doi: 10.1038/s41587-020-0472-9.

P. A. P. Moran. Notes on continuous stochastic phenomena. Biometrika, 37(1-2):17–23, June 1950. ISSN 0006-3444. doi: 10.1093/biomet/37.1-2.17.

Lambda Moses and Lior Pachter. Museum of spatial transcriptomics. Nature Methods, 19(5):534–546, May 2022. ISSN 1548-7105. doi: 10.1038/s41592-022-01409-2.

Susanne Nichterwitz, Geng Chen, Julio Aguila Benitez, Marlene Yilmaz, Helena Storvall, Ming Cao, Rickard Sandberg, Qiaolin Deng, and Eva Hedlund. Laser capture microscopy coupled with Smart-seq2 for precise spatial transcriptomic profiling. Nat Commun, 7(1):12139, July 2016. ISSN 2041-1723. doi: 10.1038/ncomms12139.

Frédéric Normandeau, Andy Ng, Maiwenn Beaugrand, and David Juncker. Spatial Bias in Antibody Microarrays May Be an Underappreciated Source of Variability. ACS Sens., 6 (5):1796–1806, May 2021. ISSN 2379-3694. doi: 10.1021/acssensors.0c02613.

Oscar E. Ospina, Alex C. Soupir, Roberto Manjarres-Betancur, Guillermo Gonzalez-Calderon, Xiaoqing Yu, and Brooke L. Fridley. Differential gene expression analysis of spatial transcriptomic experiments using spatial mixed models. Sci Rep, 14(1):10967, May 2024. ISSN 2045-2322. doi: 10.1038/s41598-024-61758-0.

Jasmine T. Plummer, Felipe Segato Dezem, David P. Cook, Jiwoon Park, Luke Zhang, Yutian Liu, Maycon Marção, Hannah DuBose, Arjumand Wani, Kellie Wise, Michael Roach, Kate Harvey, Taopeng Wang, Kirk B. Jensen, Natalia Morosini, Roberto De Gregorio, Alicia Alonso, Shauna Lee Houlihan, Robert E. Schwartz, Erika Hissong, Catherine Snopkowski, Jeffrey L. Wrana, Natalie Ryan, Lisa M. Butler, George Church, Alexander Swarbrick, Christopher E. Mason, and Luciano G. Martelotto. Standardized metrics for assessment and reproducibility of imaging-based spatial transcriptomics datasets. Nat Biotechnol, 44(7):1213–1225, July 2026. ISSN 1546-1696. doi: 10.1038/s41587-025-02811-9.

Maria Renelt, Viola von Bohlen und Halbach, and Oliver von Bohlen und Halbach. Distribution of PCP4 protein in the forebrain of adult mice. Acta Histochemica, 116(6):1056–1061, July 2014. ISSN 0065-1281. doi: 10.1016/j.acthis.2014.04.012.

Abhyudai Singh and Pavol Bokes. Consequences of mRNA Transport on Stochastic Variability in Protein Levels. Biophysical Journal, 103(5):1087–1096, September 2012. ISSN 00063495. doi: 10.1016/j.bpj.2012.07.015.

Michael A. Sirover. New nuclear functions of the glycolytic protein, glyceraldehyde-3phosphate dehydrogenase, in mammalian cells. Journal of Cellular Biochemistry, 95(1):45–52, May 2005. ISSN 0730-2312. doi: 10.1002/jcb.20399.

Shiquan Sun, Jiaqiang Zhu, and Xiang Zhou. Statistical analysis of spatial expression patterns for spatially resolved transcriptomic studies. Nat Methods, 17(2):193–200, February 2020. ISSN 1548-7105. doi: 10.1038/s41592-019-0701-7.

Valentine Svensson, Sarah A Teichmann, and Oliver Stegle. SpatialDE: Identification of spatially variable genes. Nature Methods, 15(5):343–346, May 2018. ISSN 1548-7105. doi: 10.1038/nmeth.4636.

Art Taychameekiatchai, Xiaowei Zhan, Guanghua Xiao, and Peifeng Ruan. GPSNorm: Gaussian Process Spatial Normalization for Spatial Transcriptomics, July 2026. ISSN 2692-8205.

Lukas M. Weber, Arkajyoti Saha, Abhirup Datta, Kasper D. Hansen, and Stephanie C. Hicks. nnSVG for the scalable identification of spatially variable genes using nearest-neighbor Gaussian processes. Nat Commun, 14(1):4059, July 2023. ISSN 2041-1723. doi: 10.1038/s41467-023-39748-z.

Guanao Yan, Shuo Harper Hua, and Jingyi Jessica Li. Categorization of 34 computational methods to detect spatially variable genes from spatially resolved transcriptomics data. Nat Commun, 16(1):1141, January 2025. ISSN 2041-1723. doi: 10.1038/s41467-025-56080-w.

