## Supplementary Figures for "Biological and technical spatial variation in spatial transcriptomics"

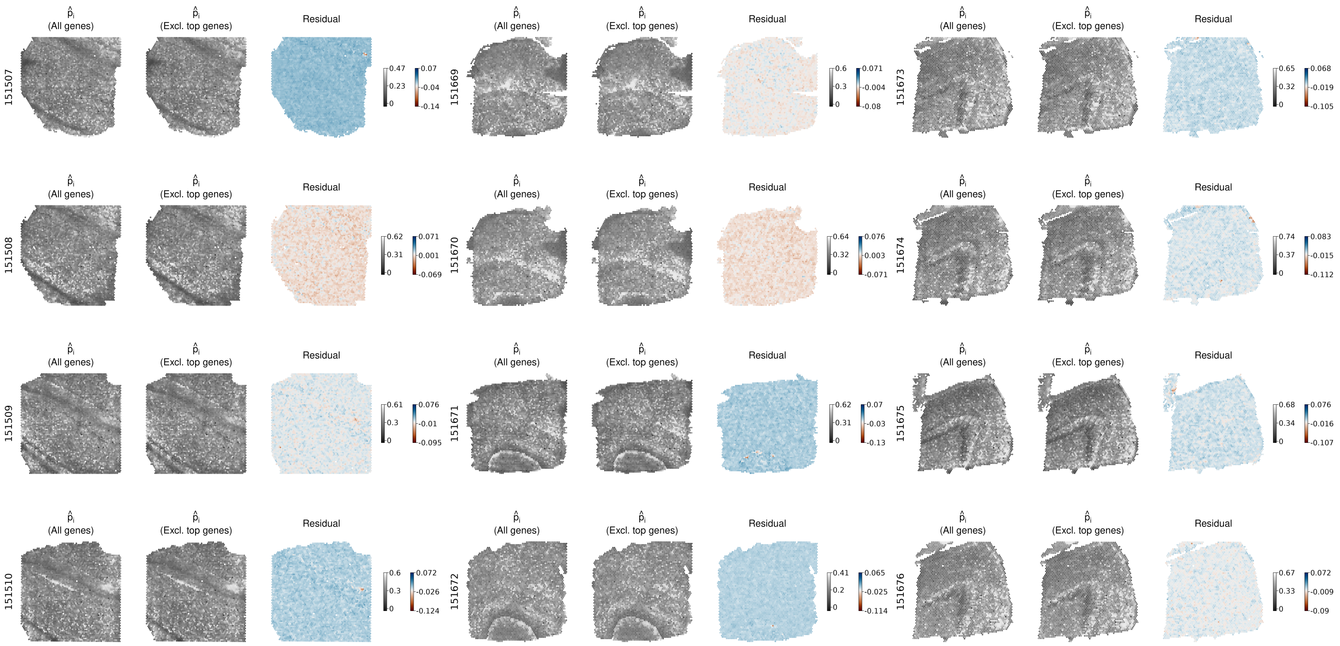


**Fig. S1.** To assess whether the inferred spatial field was driven by a small number of highly expressed genes, we re-estimated the field after removing the 25 most highly expressed genes. The inferred spatial field was consistent when estimated from the smaller subset of genes, suggesting that it reflects a shared source of variation rather than patterns driven by individual genes or gene sets.

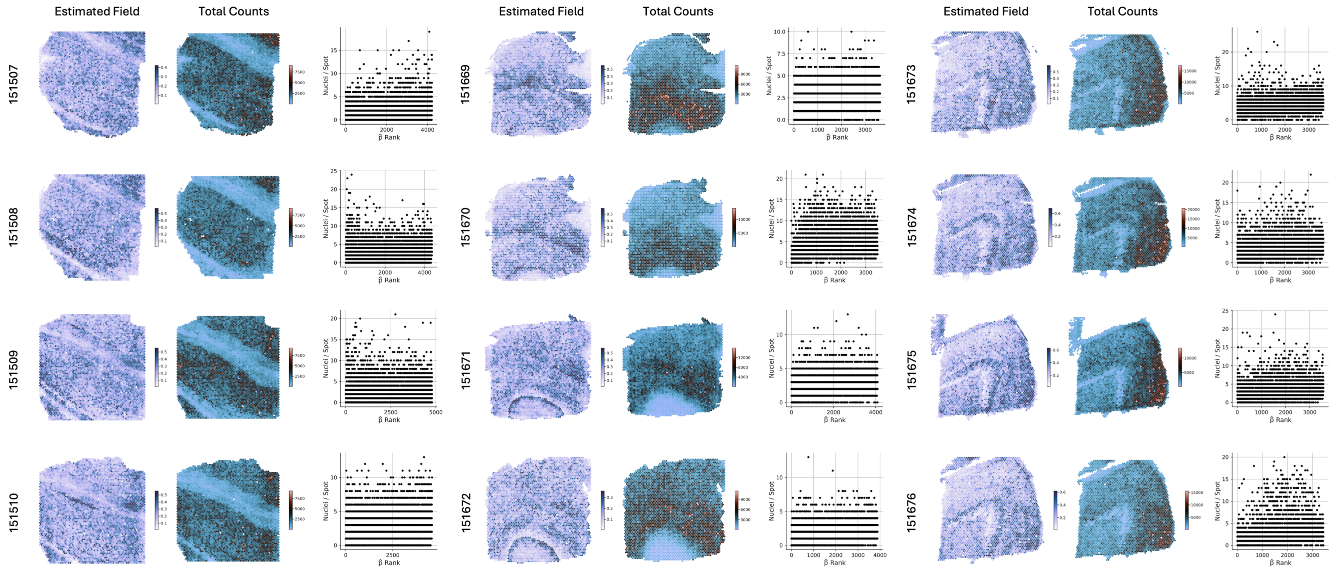


**Fig. S2.** For each of 12 DLPFC tissue sections, the estimated spatial capture field (left) is compared with the observed total transcript count at each capture location (center). Scatterplots (right) show the relationship between the rank of the estimated field at each location and the number of nuclei overlapping each spot.

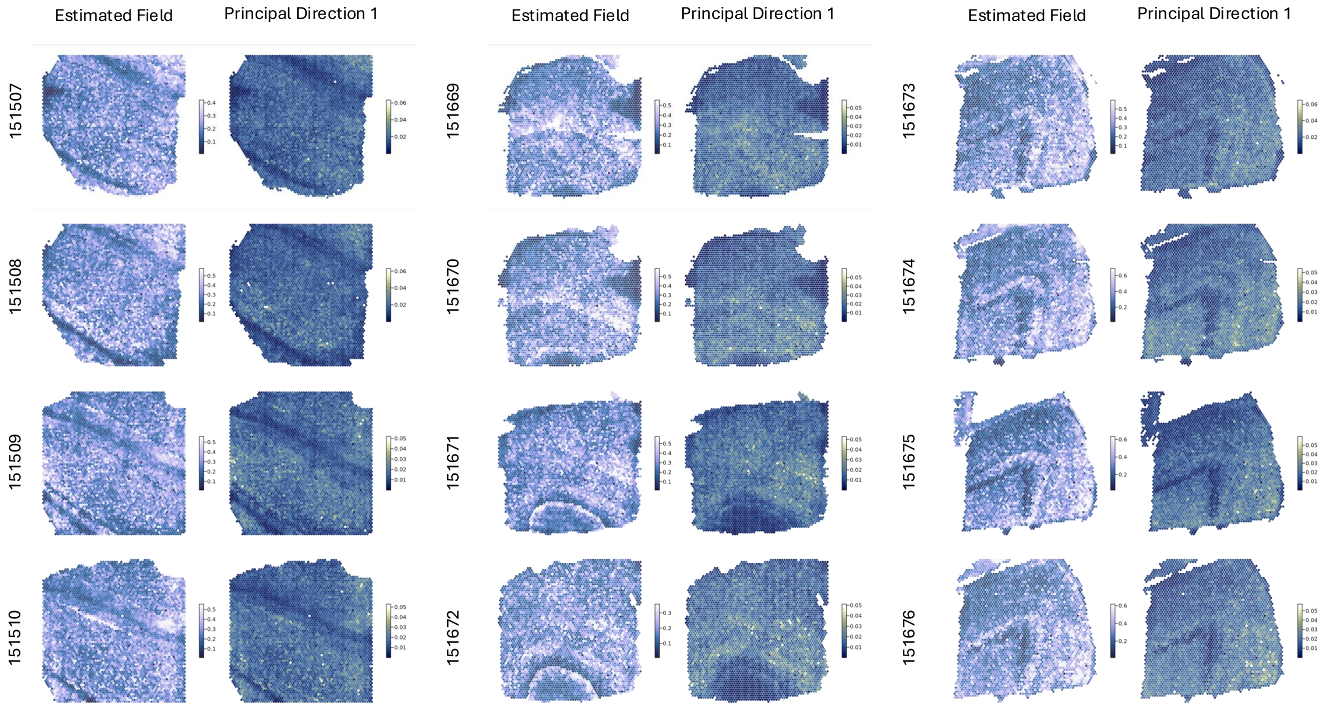


**Fig. S3.** Comparison of the estimate spatial capture field and the leading SVD component. SVD was performed on the matrix of raw residuals computed by subtracting the raw gene counts from the grouped means.

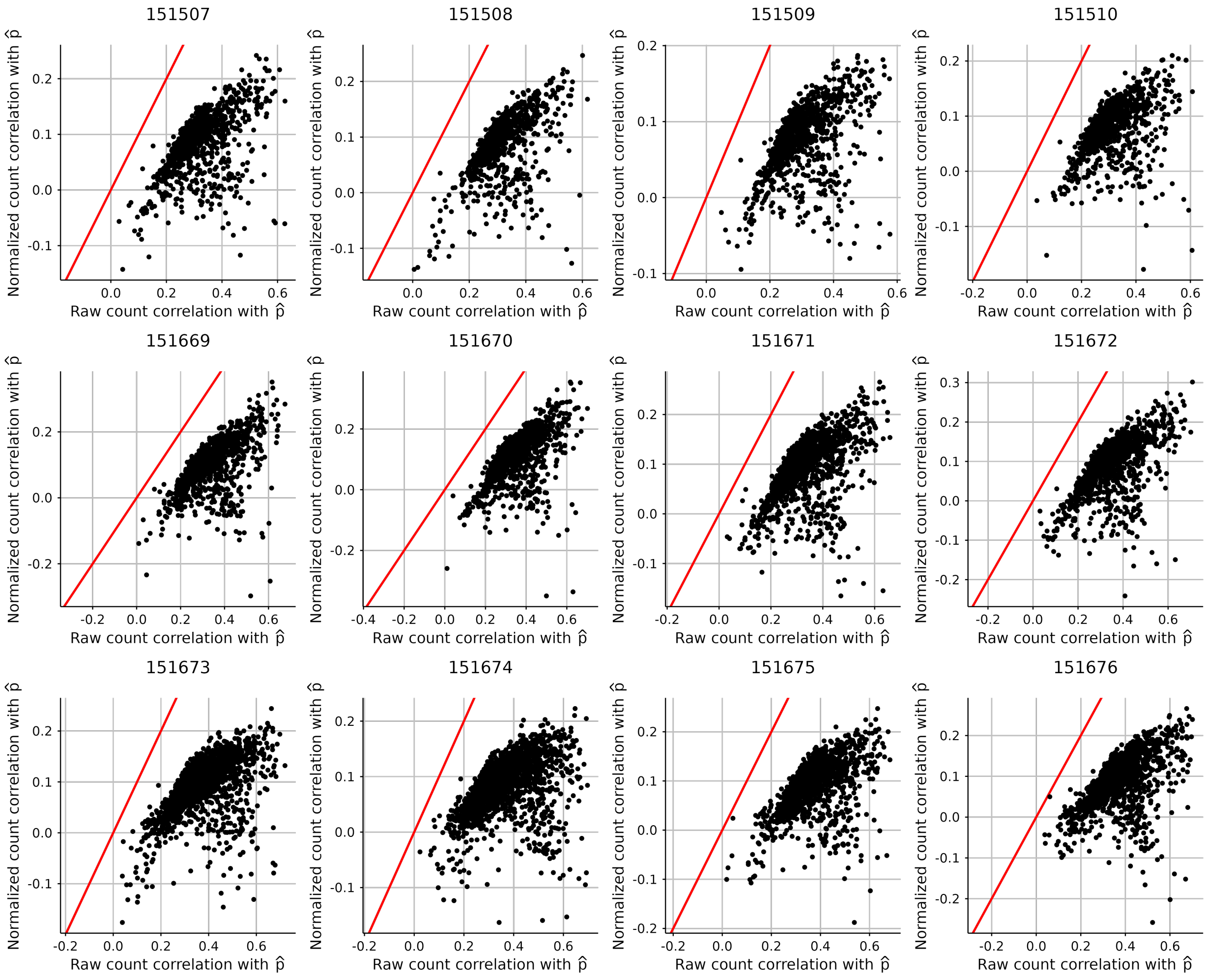


**Fig. S4.** Total-count normalization removes a spatial component that is shared between gene counts and the inferred field. The observation that normalization attenuates association with $\hat{p_i}$ is consistent with $\hat{p_i}$ representing a shared multiplicative component of the measurement process.

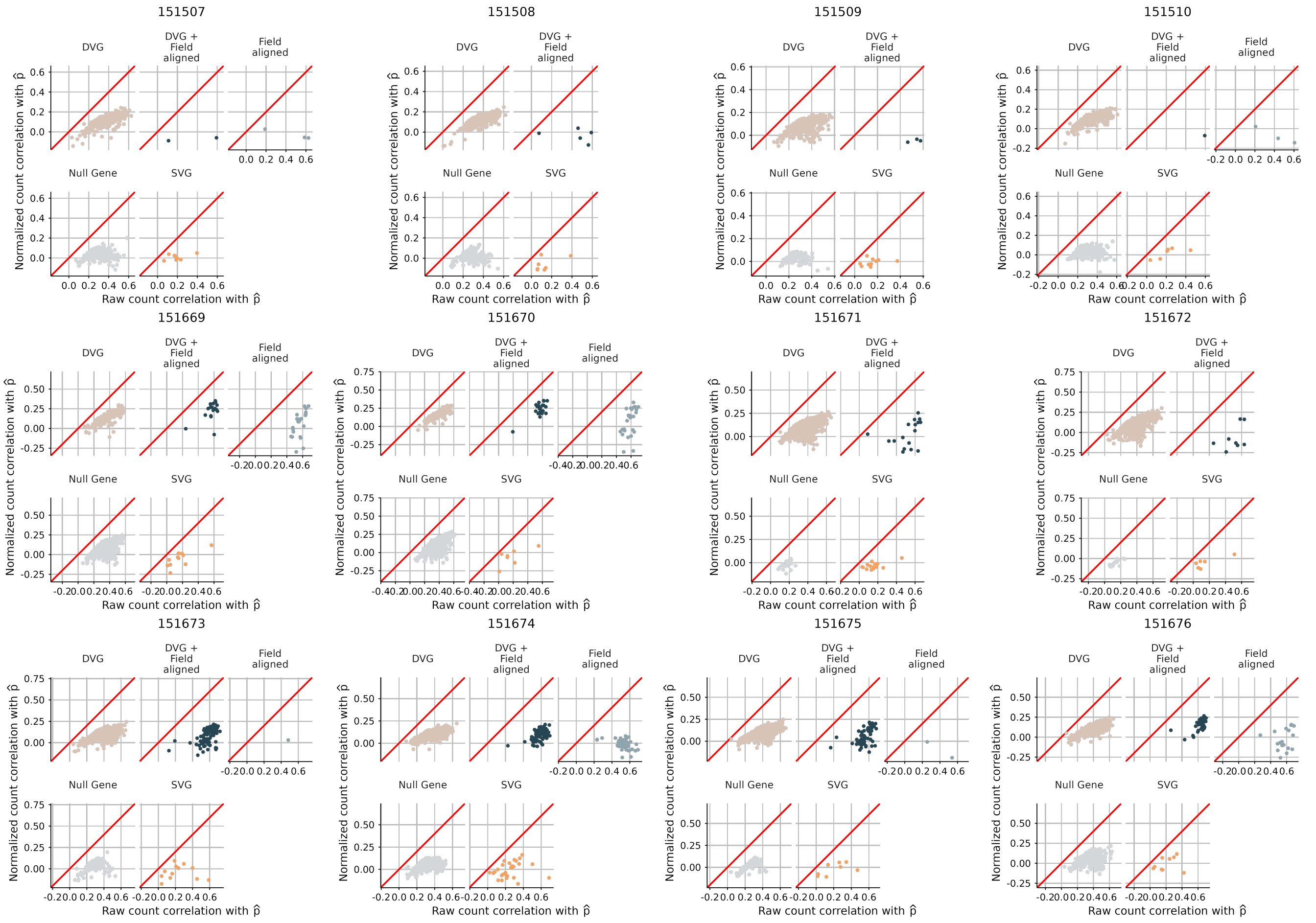

**Fig. S5.** Genes whose spatial variation is aligned with the inferred field lose more association with the field after conventional library-size normalization, providing further support for the idea that the inferred field does not capture a gene-specific spatial expression signal.

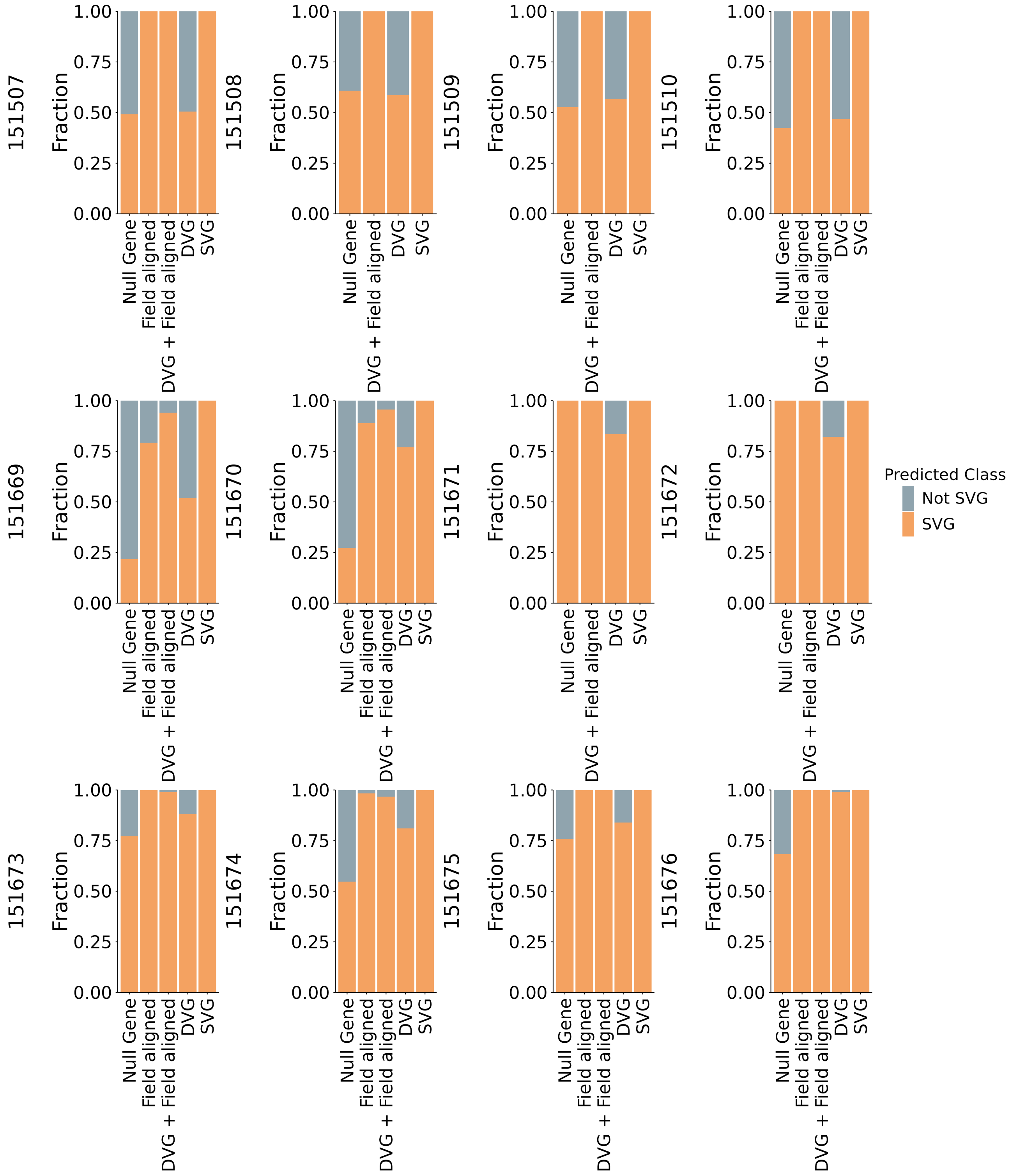


**Fig. S6.** Comparisons of gene calls made by stipple and SpatialDE. The stipple gene class is shown on the x-axis.

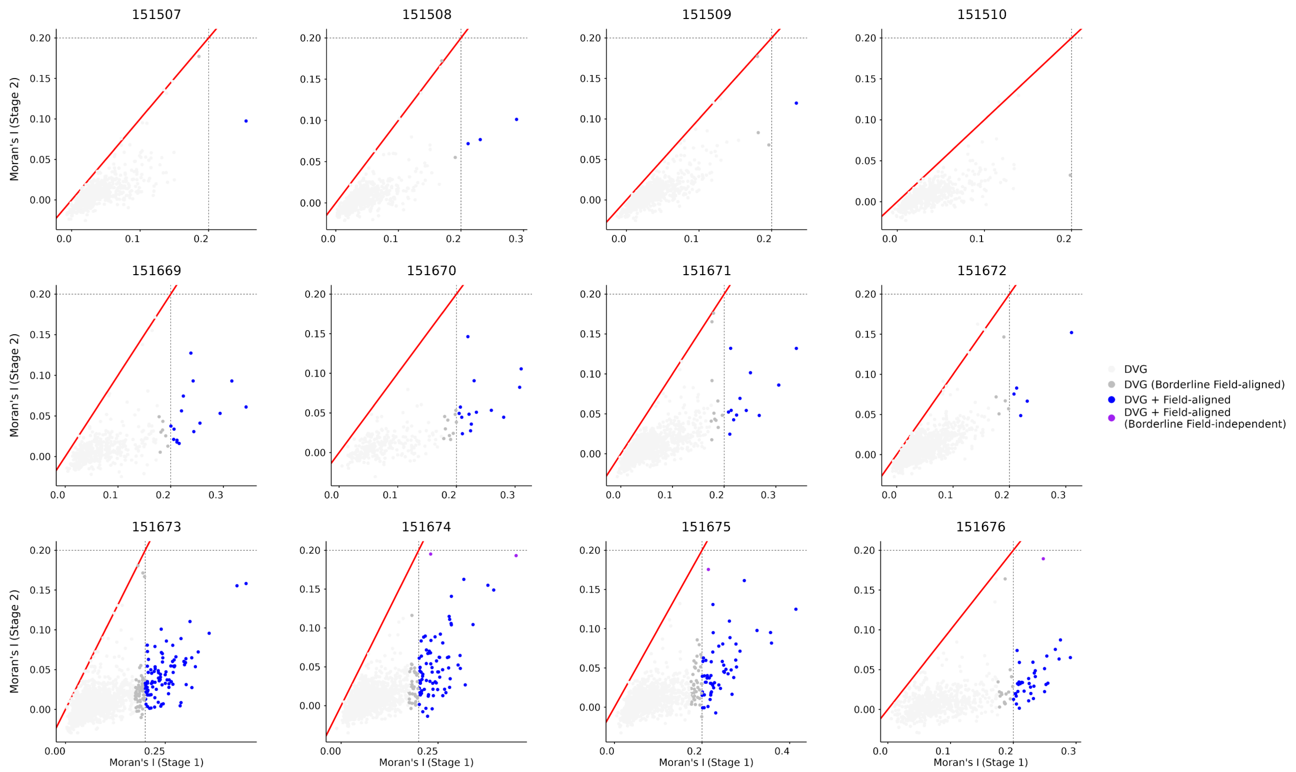


**Fig. S7.** Two-stage gene calling for each of the 12 samples in the DLPFC dataset. Genes colored in dark gray are near the borderline of the Moran's I threshold and could plausibly be labelled field-aligned.

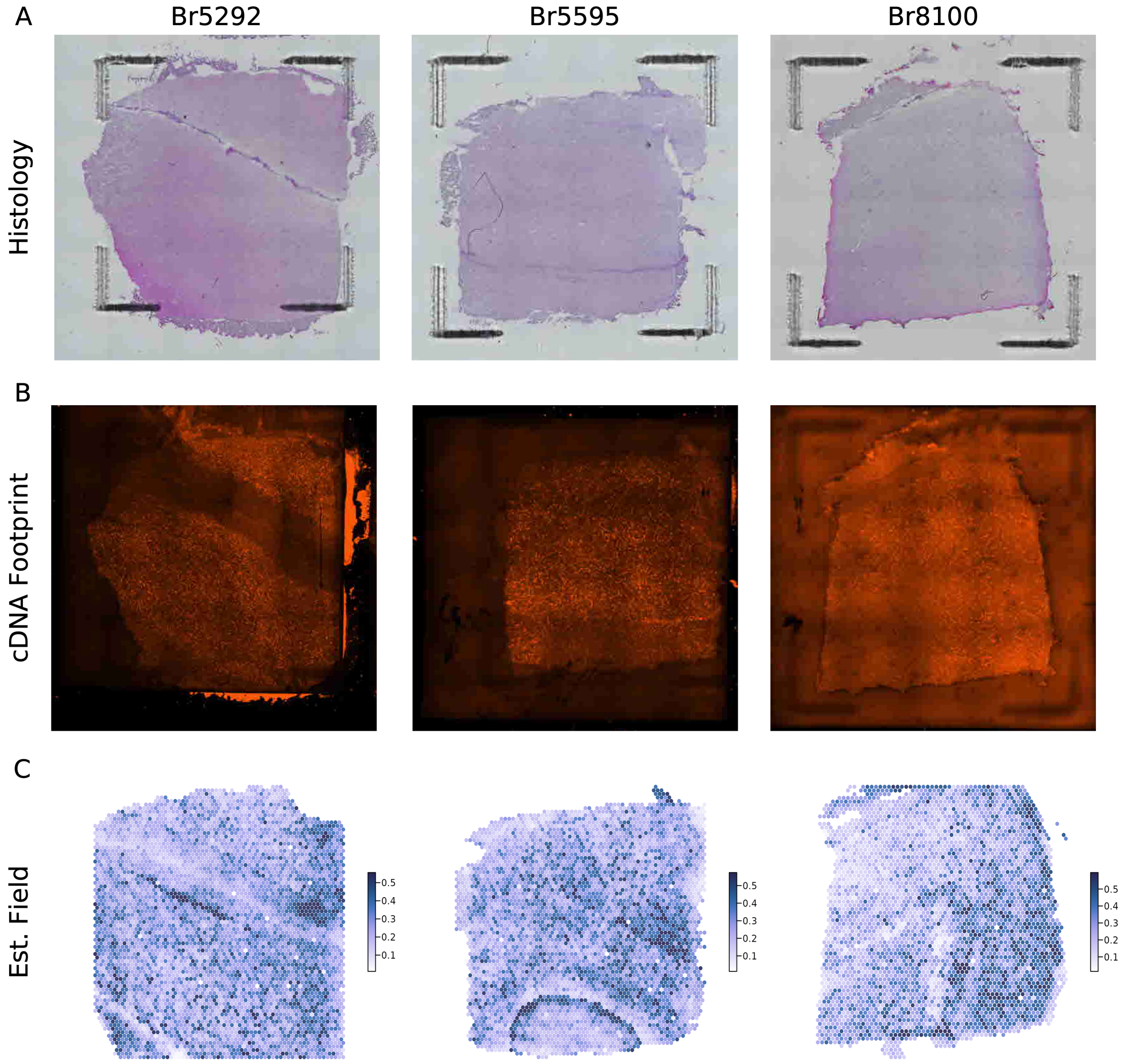


**Fig. S8.** A-B.) Recreated Supp. Fig. 1 from Maynard, et.al. (2021) showing the bright field histology images and fluorescent cDNA staining. C.) Representative spatial capture field estimates from each sample.
